# Atypical low-frequency and high-frequency neural entrainment to rhythmic audiovisual speech in adults with dyslexia

**DOI:** 10.1101/2025.09.14.675562

**Authors:** Mahmoud Keshavarzi, Brian C J Moore, Usha Goswami

## Abstract

Developmental dyslexia has been linked to atypical neural processing of the temporal dynamics of speech, but there has been disagreement concerning whether faster or slower dynamics are impaired. According to the Temporal Sampling (TS) theory, dyslexia arises from impaired entrainment of low-frequency neural oscillations – particularly in the delta (1–4 Hz) and theta (4–8 Hz) bands – to the rhythmic modulations of speech. This hypothesis was tested for adults with and without dyslexia using electroencephalography (EEG) during a rhythmic audiovisual speech paradigm, previously delivered to children. Participants viewed a “talking head” repeating the syllable “ba” at a 2-Hz rate. Measures were neural phase entrainment and band power in the delta, theta, beta (15–25 Hz), and low gamma (25–40 Hz) bands, and broadband event-related potentials (ERPs), focusing on the N1 and P2 components. Phase-amplitude coupling (PAC) and phase-phase coupling (PPC) were also assessed for delta-theta, delta-beta, theta-beta, delta-gamma, and theta-gamma interactions. Both groups exhibited significant delta- and theta-band phase entrainment; however, the two groups differed significantly in the preferred phase for the theta band. While the control group showed consistent beta- and low gamma-band phase entrainment, this was not observed for the dyslexic group. There was significantly greater delta-band power for the dyslexic group across the whole brain and in the right temporal region. Additionally, the P2 ERP component differed significantly between groups. The data are interpreted with respect to TS theory.

## Introduction

Dyslexia is a developmental disorder characterised by impairments in reading, writing, and spelling, primarily attributed to a core difficulty in phonological processing (Rack, 1994; Ramus et al., 2004). This ‘phonological deficit’ is characterised by difficulties in recognising, memorising, and manipulating sounds in words, which are precursor essential skills for learning to read (Lyon et al., 2003). Affecting approximately 5-10% of the population (Roitsch and Watson, 2019), dyslexia is not associated with impaired general intelligence, reasoning, and intellectual abilities but poses substantial challenges in academic environments and daily life (Shaywitz, 1998). A central aspect of the phonological difficulties is poor phonological awareness, defined as the capacity to recognise and manipulate oral units such as syllable stress patterns, syllables, onset-rimes, and phonemes within spoken language (Goswami et al., 2021). These challenges likely arise from atypical neural processing within brain regions critical for language and auditory functions, and may involve neural oscillatory systems (Tallal & Piercy, 1973; Goswami, 2011; Lehongre et al., 2011). The Temporal Sampling (TS) theory (Goswami, 2011, 2015, 2018) proposes that dyslexia is characterised by deficits in accurately processing slower temporal characteristics of speech sounds, with frequencies below 10 Hz, primarily affecting the delta and theta oscillatory networks. In contrast, other theories focus on rapid processing deficits in temporal windows of about 40 ms, corresponding to frequencies above 25 Hz (gamma band oscillatory networks, Tallal, 2004; Lehongre et al., 2011). Specifically, TS theory suggests that individuals with dyslexia struggle to perceive and process the precise timing and patterns of amplitude modulation associated with syllabic structures, the parsing of which is related to slower rates of temporal modulation. Tallal’s Rapid Auditory Processing (RAP) theory and the ‘phonemic oversampling’ proposed by Lehongre et al. (2011) focus on phoneme-level structures, which are related to faster rates of temporal modulation. However, given the mechanistic hierarchical nesting between slow (theta) and fast (gamma) oscillatory rates in auditory cortex (Gross et al., 2013), it is possible that atypical low-frequency temporal sampling may subsequently result in altered fast-rate activity. Any level of impaired temporal processing is assumed to hinder the development of phonological awareness in infancy and childhood, complicating subsequent learning of the mapping between written letters and their corresponding sounds.

The current study is a test of TS theory, specifically of the proposal that if the core phonological difficulties in dyslexia measured in childhood arise from atypical mechanisms of low-frequency (<10 Hz) neural rhythmic entrainment during speech processing, then these mechanisms should still be atypical in adulthood. Prior studies of neural entrainment utilising rhythmic inputs in adults have utilised non-speech stimuli, most typically amplitude-modulated (AM) noise (McAnally & Stein, 1997; Menell et al., 1999; Lehongre et al., 2011; Poelmans et al., 2012). However, they have either not measured oscillations in the delta band (∼2 Hz, Poelmans et al., 2012), or have not measured oscillations in both the delta and theta bands (modulations < 10Hz, McAnally & Stein, 1997; Menell et al., 1999; Lehongre et al., 2011). Using AM white noise, McAnally and Stein (1997) reported reduced Auditory Steady State Response (ASSR) power to AM with rates from 20 to 80 Hz in dyslexic adults compared to controls. Menell et al. (1999) measured ASSR power for AM rates of 10, 20, 40, 80 and 160 Hz and reported that the ASSR power was weaker for dyslexic readers than for controls for all AM rates. However, both these studies used only a single electrode. Lehongre et al. (2011) used a white noise stimulus with a range of AM rates (10–80 Hz), which they argued broadly covered the phonemic sampling domain. Using magnetoencephalography (MEG), they demonstrated lower left-lateralised low-gamma (∼30 Hz) responses in dyslexic adults than for typically-reading control adults, which was interpreted to show impaired entrainment to phonemic-rate modulations in left auditory cortex. Intriguingly, they also observed enhanced cortical entrainment at rates >40 Hz in the dyslexic adults, which they reported was linked to impaired phonological memory. Using EEG, Poelmans et al. (2012) administered speech-weighted noise amplitude modulated at 4, 20, and 80 Hz to measure ASSRs in dyslexic adults. They employed a limited EEG montage (recordings from four electrodes). They reported no differences between dyslexic and control adults at 4 Hz or at 80 Hz, but a significant dyslexic deficit at 20 Hz. Poelmans et al. (2012) concluded that cortical processing of ‘phonemic-rate’ auditory modulations was selectively impaired in adults with dyslexia. As a test of TS theory, Hämäläinen et al. (2012) played AM white noise at 4 rates (2, 4, 10, and 20 Hz) to adults with and without dyslexia. They reported significantly reduced phase entrainment for the dyslexic participants in right hemisphere networks, but for the 2-Hz rate only. They also found significantly weaker right hemisphere entrainment overall (summed across modulation rates) for the dyslexic participants, and significantly stronger entrainment to the 10-Hz rate in the left hemisphere, a finding that was not predicted. This could indicate compensatory entrainment at faster temporal rates in those with dyslexia, consistent with the finding reported (bilaterally) by Lehongre et al. (2011) for rates above 40 Hz.

One possible reason for the discrepant findings across these studies using nonspeech stimuli is that participants had learned to read different languages, French (Lehongre et al., 2011), Dutch (Poelmans et al., 2012) and English (McAnally & Stein, 1997; Menell et al., 1999; Hämäläinen et al., 2012). However, this body of work has also been critiqued for the reliance on non-speech stimuli: the atypical patterns of entrainment may lack linguistic relevance. For the EEG paradigms used by McAnally & Stein (1997), Menell et al., (1999) and Poelmans et al. (2012), low spatial resolution is a further issue (due to the sparse electrode montage). Accordingly, it is important to examine neural entrainment to rhythmic speech rather than non-speech stimuli, which may have greater ecological validity and clinical relevance. To address this gap in the adult literature, here we utilised a rhythmic syllable repetition task previously used with dyslexic children (Power et al., 2013; Keshavarzi et al., 2022) and infants (Ni Choisdealbha et al., 2023). The current study investigated responses in the beta and low gamma bands in addition to the delta and theta bands, in order to investigate whether atypical entrainment for dyslexic adults would be found at 20 Hz (Poelmans et al., 2012) or near 35 Hz (Lehongre et al., 2011) for a rhythmic speech input.

Each of oscillatory band is thought to serve a specific role in the neural processing of speech. Delta and theta oscillations are involved in processing the rhythmic characteristics of speech and are particularly important for phrasal (delta) and syllabic (theta) processing (Giraud & Poeppel, 2012). Infant EEG research suggests that these low-frequency bands are the foundational components necessary for the development of an efficient language system (Attaheri et al., 2024; Keshavarzi et al., 2024a), as infants show significant entrainment to natural speech in the delta and theta but not alpha bands. In infant longitudinal EEG studies, individual differences from age 2 months in preferred phase in the rhythmic syllable repetition task were found to predict differences in vocabulary and phonology at age 2 years (Ni Choisdealbha et al., 2023). Beta oscillations are implicated in motor functions (Engel & Fries, 2010) and play a significant role in top-down attentional regulation and sensory event prediction (Arnal et al., 2015). In the music domain, beta-band oscillations have been thought to play a key role in representing auditory beats, that is representing the temporal structure of regular sound sequences (Fujioka et al., 2012, 2015). Low-gamma oscillations are thought necessary for encoding the rapid acoustic information necessary for phonemic representation (Lehongre et al., 2011; Giraud & Poeppel, 2012). These fast oscillations may support encoding of the fine-grained temporal details essential for distinguishing rapid transitions in speech, such as formant transitions. Disruptions in gamma-band activity have been observed in individuals with dyslexia (Lehongre et al., 2011). Lehongre et al. (2011) interpreted atypical gamma band entrainment to AM noise as a neural explanation for dyslexic difficulties in phoneme-level processing.

Neural phase entrainment has been studied intensively in dyslexia, as it is one of the mechanisms that enables the brain to align temporally with external rhythmic stimuli, thus facilitating efficient processing of rhythmic information (Lakatos et al., 2008). In the rhythmic syllable repetition task, children with dyslexia exhibit atypical phase entrainment in both the delta and beta frequency bands (Power et al., 2013; Keshavarzi et al., 2022, 2024b). Additionally, numerous studies that have investigated ERPs for dyslexic individuals have found changes in the latency and amplitude of ERPs. For example, several studies with both children and adults reported abnormal early ERP components, such as N1 and P2, which are associated with the initial processing of auditory stimuli (Bernal et al., 2000; Helenius et al., 2002; Hämäläinen et al., 2007; Peter et al., 2019). ERPs represent time-locked electrophysiological responses directly related to specific sensory, cognitive, or motor events. Alterations in ERP components may reflect delayed or atypical sensory processing (e.g., auditory and visual) in dyslexic individuals, suggesting underlying neural inefficiencies.

Other studies have reported abnormal neural oscillatory band-power in children with dyslexia (Arns et al., 2007; Cutini et al., 2016; Power et al., 2016; Keshavarzi et al., 2024b). Band-power analyses quantify neural oscillation power within distinct frequency ranges, providing insights into the strength of neural responses. Another common neural measure in speech processing studies is cross-frequency coupling, encompassing phase-amplitude coupling (PAC) and phase-phase coupling (PPC, e.g. Giraud & Poeppel, 2012; Gross et al., 2013). PAC and PPC have not to date been explored in adult dyslexia studies, to our knowledge. PAC refers to the interaction where the phase of a lower-frequency oscillation is related to the amplitude of a higher-frequency oscillation (Tort et al., 2008). PAC may facilitate communication between neural networks operating in different frequency bands (Canolty and Knight, 2010). Abnormalities in cross frequency coupling, particularly theta-gamma PAC, are believed to disrupt the timing and coordination of the neural activity necessary for phonological processing and reading fluency (Archer et al., 2020). In infant longitudinal studies, infants with stronger theta-gamma PAC at age 4 months showed better language outcomes at age 2 years (Attaheri et al., 2024). PAC is a currently neglected aspect of neural temporal dynamics in the adult dyslexia literature, and accordingly was measured here. PPC is a measure of the temporal synchrony between oscillations in different frequency bands (Tass et al., 1998; Lachaux et al., 1999) and is also lacking in current adult dyslexia studies. In a speech context, differences in PPC could reflect impairments in the timing and coordination of neural processes that are important for efficient auditory and phonological processing. PPC is also computed here.

In summary, in the current study a rhythmic audio-visual task (Power et al., 2013; Keshavarzi et al., 2022, 2024c) was administered to 24 adults with dyslexia and 24 adults without dyslexia while concurrently recording EEG data. We examined phase entrainment and neural oscillatory band-power within the delta (0.5–4 Hz), theta (4–8 Hz), beta (15–25 Hz), and low gamma (25–40 Hz) frequency bands. Additionally, broadband ERPs were computed separately for each group, focusing on the N1 and P2 components. Furthermore, PPC and PAC were assessed between delta-theta, delta-beta, and theta-beta frequency pairs for each group independently. Consistent with our previous findings using the syllable repetition paradigm with dyslexic children (Keshavarzi et al., 2024b), we hypothesised *a priori* that dyslexic group would exhibit an absence of consistent neural phase entrainment in the beta and gamma frequency bands. Based on TS theory and previous findings from Power et al. (2013) and Keshavarzi et al. (2022), we predicted that the preferred entrainment phase in both delta and theta bands would differ significantly between dyslexic and control groups. Consistent with existing evidence (Bernal et al., 2000; Helenius et al., 2002; Hämäläinen et al., 2007; Peter et al., 2019), we expected that dyslexic individuals would demonstrate atypical ERP responses, specifically abnormal amplitudes of N1 and P2 components. We further predicted significantly elevated band-power for the dyslexic group in the delta frequency band both across the entire brain and within the right temporal region (Arns et al., 2007; Cutini et al., 2016), as well as in the beta frequency band across the whole brain (Power et al., 2016; Keshavarzi et al., 2024b). No significant group differences were anticipated for delta-beta PAC, as to date no differences have been found in studies of children with dyslexia (Power et al., 2016; Keshavarzi et al., 2024b). We did not have specific hypotheses regarding group differences in PAC for delta-theta, theta-beta, delta-low gamma, and theta-low gamma bands, or in PPC for delta-theta, delta-beta, theta-beta, delta-low gamma, and theta-low gamma frequency pairs.

## 2. Methods and Material

### 2.1. Participants

A total of 48 adult native English speakers were tested, comprising 24 participants diagnosed with dyslexia (mean age = 21.4, standard deviation (SD) ± 2.5 years, age range = 18.3–29.1 years) and 24 control participants without dyslexia (mean age = 21.9, SD = 2.6 years, age range = 18.7–29.2 years). Dyslexic participants were recruited via the Accessibility and Disability Resource Centre at the University of Cambridge and had received an official diagnosis of dyslexia. The inclusion criteria required that dyslexic individuals had no comorbid learning disabilities, such as autism spectrum disorder, dyspraxia, attention-deficit/hyperactivity disorder, or developmental language disorder, as verified through participant self-report. Additionally, dyslexic participants were required to perform at least one standard deviation below the control group mean on at least one subtest of the Test of Word Reading Efficiency, Second Edition (TOWRE-2; Torgesen et al., 2012). Control participants were matched closely to the dyslexic group in terms of age and intellectual functioning. All participants underwent audiometric screening, assessing pure-tone auditory sensitivity bilaterally at frequencies of 250, 500, 1000, 2000, 4000, and 8000 Hz. Participants were required to have auditory thresholds of 20 dB HL or better. All participants provided written informed consent prior to their involvement, consistent with ethical guidelines set forth by the Declaration of Helsinki. The study was reviewed by the Psychology Research Ethics Committee at the University of Cambridge and received a favourable opinion.

### 2.2. Experimental set-up and stimuli

The experimental design and stimulus presentation followed the procedures described in Keshavarzi et al. (2022, 2024c). Following EEG cap placement, participants were seated in an electrically shielded, soundproof room. They heard and watched audio-visual stimuli via earphones and on a screen, respectively. EEG signals were recorded at a sampling rate of 1 kHz using a 128-channel EEG system (HydroCel Geodesic Sensor Net). Stimuli consisted of a video display of a female talker producing rhythmic sequences of the syllable “ba” at a 2 Hz rate. Each sequence comprised 14 syllables, and the face of the talker was presented 68 ms before the onset of each syllable. One of the syllables in each sequence (either the 9th, 10th, or 11th) was temporally shifted to create a rhythmic violation. Participants were instructed to focus on the speaker’s lips while attending to the auditory stream and to press a key when they detected a syllable presented out of rhythm. The shift varied across trials. A 3-down 1-up adaptive staircase procedure was employed to determine the shift that was just noticeable for each participant. When a participant correctly identified the rhythmic violations on three consecutive trials, the deviation from the isochronous 500-ms stimulus-onset asynchrony (SOA) was reduced by 16.67 ms. A single failure to detect a violation led to an increase in the SOA of 16.67 ms. This procedure was designed to track an accuracy of 79.4% (Levitt, 1971), ensuring consistent task difficulty across participants. Each participant completed 90 trials, including 15 randomly interspersed catch trials containing no rhythmic deviation. Each trial was segmented into three distinct phases: an entrainment period, a violation period, and a return-to-isochrony period (see Figure 1). The EEG recording session, excluding setup time, lasted approximately 15 minutes.

**Figure 1.**
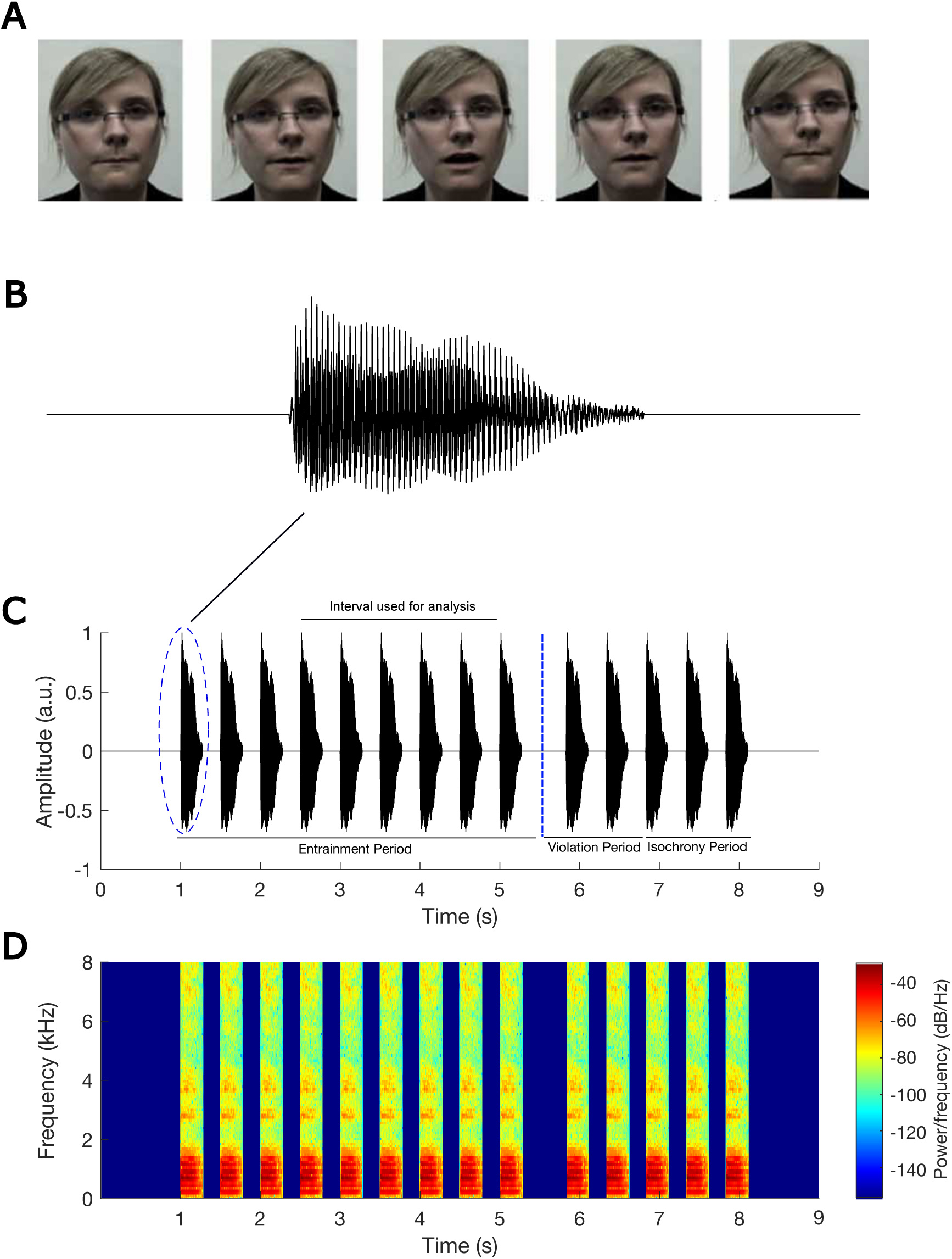
Experimental stimuli for the audiovisual task. Panel A shows a series of visual frames corresponding to a single articulation of the syllable “ba”. Panel B presents the acoustic waveform of one “ba” syllable. Panel C depicts a representative trial comprising a sequence of 14 “ba” stimuli, with a rhythmic deviation introduced at the 10th position. Panel D illustrates the corresponding spectrogram for the auditory sequence shown in Panel C. This figure is reproduced with permission from Keshavarzi et al. (2022).

### 2.3. EEG data pre-processing and analyses

The EEG recordings were initially re-referenced to the Cz channel and subsequently subjected to band-pass filtering (0.2–42 Hz) using a zero-phase finite impulse response filter with 6-dB down points at 0.1 Hz and 47.25 Hz, as implemented through the MNE-Python package (Gramfort et al., 2013). Bad channels were detected manually from inspection of the spectrum and time domain representations of the channel signals, followed by spatial interpolation (MNE-Python). Independent Component Analysis (ICA) employing the Infomax algorithm (MNE-Python) was performed on individual participant datasets. The resulting independent components were manually evaluated to remove artefactual components (e.g. eye blinks, eye movements, and heartbeat). This evaluation was based on inspection of the scalp map, power spectrum, and time-domain waveform of the components. Following preprocessing procedures, the EEG data were downsampled to 200 Hz and filtered into delta (0.5–4 Hz), theta (4–8 Hz), beta (15–25 Hz), and low gamma (25–40 Hz) bands (using MNE-Python). This filtering step was not employed for ERP and power analyses. The data were then epoched into individual trials, from 0.5 seconds before the onset of the first “ba” stimulus to 4 seconds after. To reduce computational costs, the data were downsampled to 100 Hz. The analyses were restricted to EEG responses corresponding to the 4th-8th “ba” stimuli within the entrainment period of each trial, ensuring that the rhythmicity had established.

### 2.4. Computation of phase entrainment

To evaluate phase entrainment for each group across frequency bands, the following procedure was applied, following Keshavarzi et al. (2022, 2024c):

1. The instantaneous phase for each EEG channel was computed at the time points corresponding to the onsets of the five target “ba” stimuli (4th–8th) within each trial. The instantaneous phase *φ*(*t*) of a signal *s*(*t*) was calculated using the Hilbert transform, as follows:

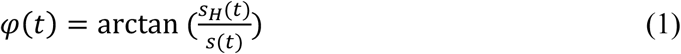

where *S_H_*(*t*) is the Hilbert transform of signal *s*(*t*), defined by:

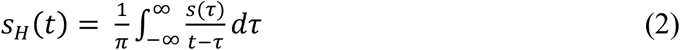
2. For each trial, the mean phase was computed by averaging the instantaneous phase values across all EEG channels and the five “ba” stimuli, resulting in a single-phase value per trial.
3. For each trial, a single unit vector (magnitude = 1) was placed in the vector space, with its angle determined by the phase value obtained in Step 2.
4. The mean vector for each participant was then calculated by averaging the unit vectors from Step 3 across trials, resulting in a single vector per participant, referred to as the *individual resultant vector*. The angle of the individual resultant vector is called the *individual preferred phase*, and its length serves as a measure of phase consistency strength across trials for each participant.
5. A single unit vector (magnitude = 1) – whose angle was determined by the angle of the *individual resultant vector* – was considered in the vector space for each participant.
6. Finally, the mean vector for each group was calculated by averaging across the unit vectors from Step 5, resulting in a single vector called the *group resultant vector*. The angle of the *group resultant vector* is called the *group preferred phase*.

### 2.5. Computation of the band-power

To investigate the band-power of neural response over the time interval used for analysis (See Figure 1C), the following steps were conducted for each individual and each group:

1. The broadband power spectral density (PSD) was computed for each EEG channel using the *pspectrum*() function in MATLAB.
2. For each trial, the broadband PSD was calculated by averaging across the broadband PSDs obtained (in step 1) for all EEG channels.
3. For each participant, the broadband PSD was calculated by averaging across the broadband PSDs obtained (in step 2) for all trials of that participant.
4. The PSD for each frequency band (delta, theta, and beta) and for each participant was calculated based on the broadband PSD obtained in step 3.
5. The PSD for each group and each frequency band (delta, theta, and beta) was calculated by averaging across the PSDs obtained (in step 4) for all participants in a given group.

### 2.6. Computation of power spectral topographic map

To compute the power spectral topographic map for each frequency band of interest (delta, theta, beta), the following steps were performed:

1. To ensure data quality, outer EEG channels (E17, E48, E119, E125, E126, E127, E128) were excluded.
2. The PSD for each epoch was computed using Welch’s method across a broad frequency range (0.2–42 Hz).
3. Spectral data were averaged within each group (dyslexic or control), and mean spectral power was then calculated for the delta, theta, and beta bands.
4. Statistical comparisons were conducted to assess power differences between groups, with a specific focus on the right temporal region (EEG channels: E100, E101, E102, E107, E108, E109, E113, E114, E115, E116, E120, E121).

### 2.7. Computation of cross-frequency PAC

Cross-frequency PAC refers to the relationship between the amplitude of a high-frequency oscillation and the phase of a low-frequency oscillation. The PAC was quantified the modulation index (*MI*), as proposed by Tort et al. (2008) and Hülsemann (2019):

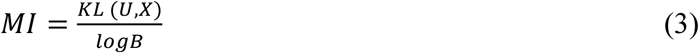

where *B* (= 18) represents the number of bins, *U* denotes the uniform distribution, *X* is the empirical distribution of the data, and *KL* (*U*, *X*) is Kullback–Leibler divergence, computed as:

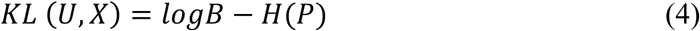

Here, *H*(.) represents the Shannon entropy, and *P* is the vector of normalised mean amplitudes per bin which is calculated as:

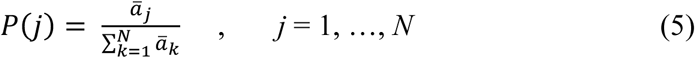

where ̄𝑎_*j*_ denotes the average amplitude within the *j*th bin, and *k* is the index over all *N* bins. Note that *P* is a vector containing *N* elements.

### 2.8. Computation of cross-frequency PPC

Cross-frequency PPC refers to the phase synchrony between oscillations in two different frequency bands. To quantify this synchrony, we used the phase locking value (PLV), as described by Lachaux et al. (1999), defined as:

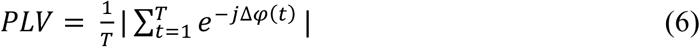

where | . | denotes to the absolute value operator, *T* represents the number of time samples, and Δ𝜑(*t*) is the phase difference, computed as:

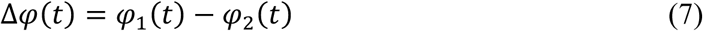

Here, 𝜑_&_(𝑡) is the instantaneous phase of first signal, and 𝜑_=_(𝑡) is the instantaneous phase of the second signal.

### 2.9. Computation of ERP

To compute the ERP waveform for each group during the entrainment period, the following steps were performed:

1. Events corresponding to each “ba” stimulus (4th–8th) were identified within the EEG timeline to ensure accurate alignment of neural responses to the stimuli.
2. Epochs were extracted around these events, ranging from 200 ms before to 400 ms after each “ba”. This time window enabled analysis of both the pre-stimulus baseline and the post-stimulus response. All epochs were baseline-corrected using the pre-stimulus interval to ensure consistent measurements across trials and participants.
3. Noisy channels within each epoch – identified by amplitudes exceeding a threshold of 60 µV – were interpolated using data from neighbouring electrodes to preserve the spatial integrity of the EEG signals.
4. The cleaned epochs were averaged for each participant to produce individual ERP waveforms.
5. Individual ERPs were then grouped and averaged for each group (control or dyslexia) to reveal potential group-level differences in neural responses to rhythmic audio-visual stimuli.
6. Statistical comparisons were conducted to assess differences in the N1 and P2 components between the two groups.

## 3. Results

### 3.1. Phase entrainment consistency for each group

To assess the phase entrainment consistency within each group and across different frequency bands, we applied the Rayleigh test of uniformity to the *individual preferred phase*s (see Section 2.4) for each group and each band (see Figure 2). For the control group, significant phase entrainment consistency was evident across all frequency bands: delta (*z* = 11.76, *p* = 1.6×10^-6^; see Figure 2A), theta (*z* = 15.66, *p* = 5.4×10^-9^; see Figure 2B), beta (*z* = 4.25, *p* = 0.013; see Figure 2C), and low gamma (*z* = 3.04, *p* = 0.046; see Figure 2D). For individuals with dyslexia, significant phase entrainment was found for the delta (*z* = 10.98, *p* = 4.53×10^-6^; see Figure 2E) and theta (*z* = 12.16, *p* = 9.46×10^-7^; see Figure 2F) bands. However, no consistent phase entrainment was observed for the beta (*z* = 1.09, *p* = 0.34; see Figure 2G) and low gamma (*z* = 1.36, *p* = 0.26; see Figure 2H) bands.

**Figure 2.**
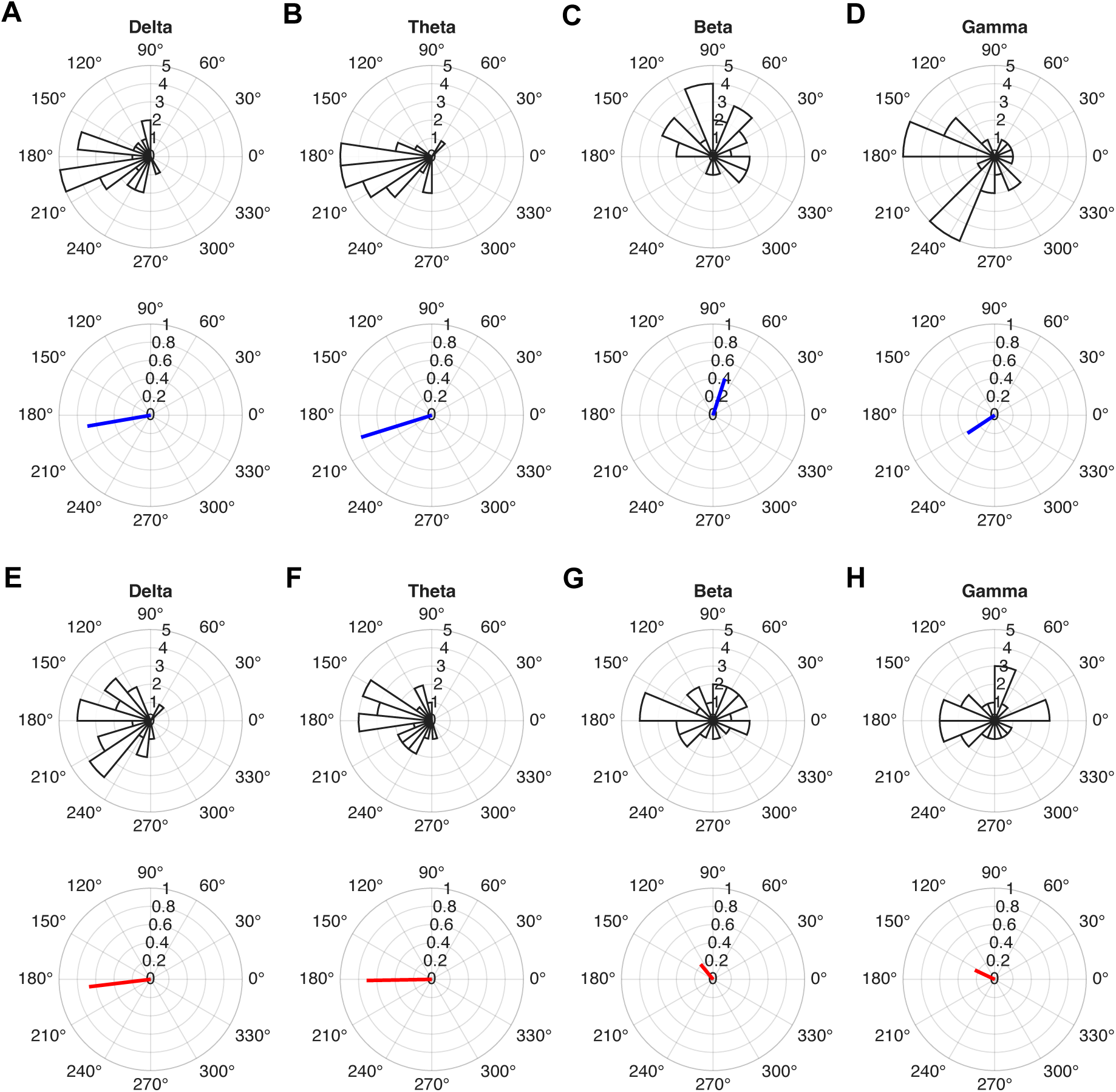
Distribution of preferred phase for each group (rose plots), along with group resultant vectors (blue and red lines). The blue vectors indicate the *group resultant vectors* for the control group, while the red vectors represent those for the dyslexic group. The control group demonstrated significant phase entrainment for the delta (Panel A), theta (Panel B), beta (Panel C), and low gamma (Panel D) bands. The dyslexic group showed significant phase entrainment for the delta (Panel E) and theta (Panel F) bands, but not for the beta (Panel G) and low gamma (Panel H) bands.

### 3.2. Comparison of preferred phase across groups

According to TS theory and to findings from Power et al. (2013) and Keshavarzi et al. (2022), we had hypothesised a group difference in preferred phase within the delta and theta bands. To test this, we conducted separate one-sample mean direction tests to compare group-level preferred phases, focusing only on frequency bands that showed significant phase entrainment. A significant difference in preferred phase was observed between groups for the theta band (*p* = 0.04), but not for the delta band (*p* = 0.77), contrary to our hypothesis.

### 3.3. Comparing the consistency of phase entrainment across groups

The length of each participant’s *individual resultant vector* (see Step 4 in Section 2.4) reflects the consistency of phase entrainment across trials, longer vectors indicating higher within-subject consistency (Keshavarzi et al., 2022). To assess group differences in phase entrainment consistency, separate two-sample t-tests were performed on the lengths of the *individual resultant vectors*, limited to frequency bands that showed significant phase entrainment (see Figure 3). No significant differences were found between the control and dyslexic groups for either the delta band (*p* = 0.33; Figure 3A) or the theta band (*p* = 0.93; Figure 3B). Phase entrainment consistency between the delta and theta bands was compared within each group. Theta-band entrainment showed significantly higher consistency than delta-band entrainment for both the control group (*p* = 0.006) and the dyslexic group (*p* = 3.1×10⁻⁵).

**Figure 3.**
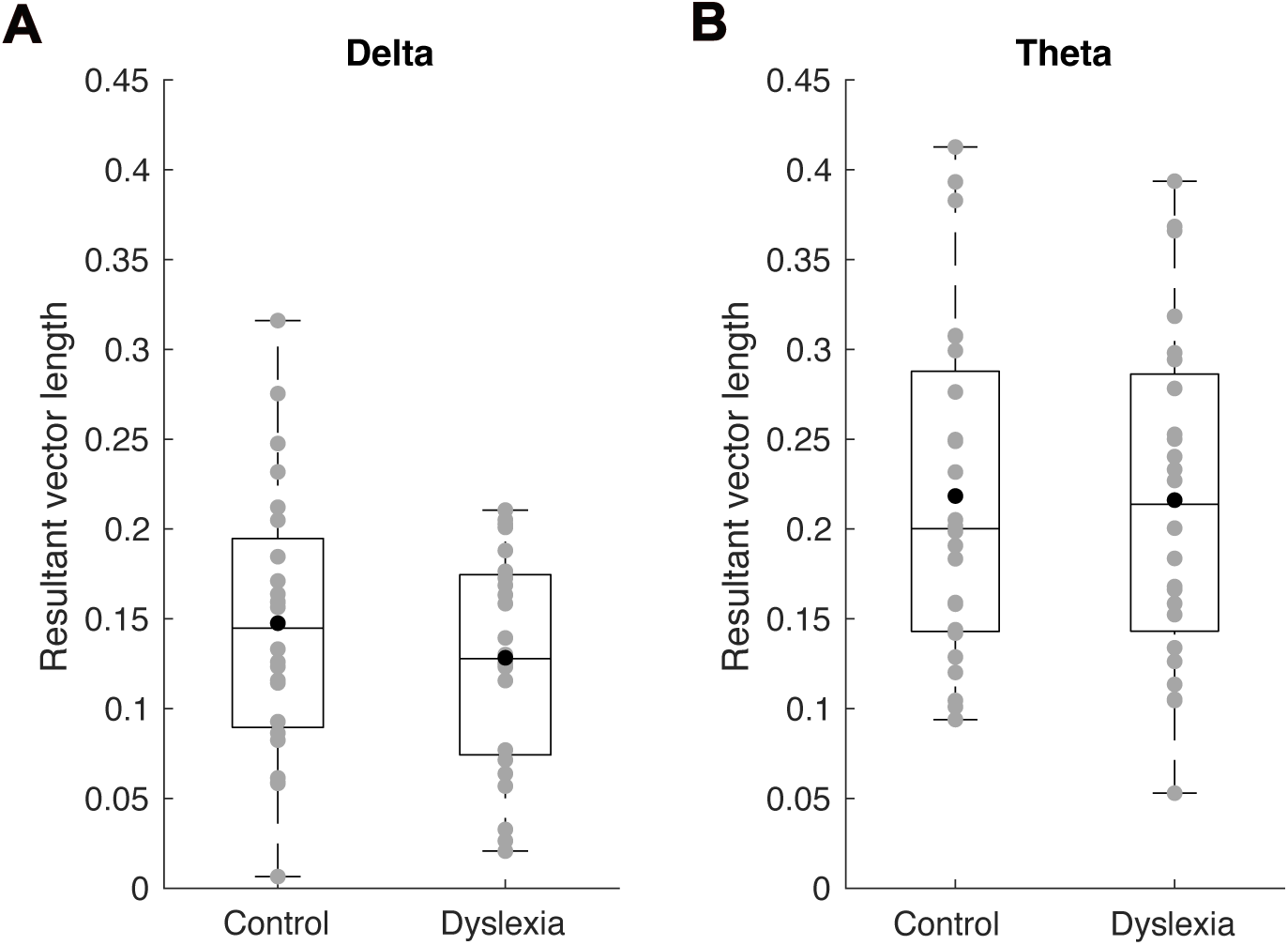
Boxplots illustrating the lengths of *individual resultant vectors*, representing the consistency of phase entrainment, for the delta (Panel A) and theta (Panel B) frequency bands. Grey circles show individual participant values, black circles indicate group means, and the horizontal line within each box marks the median. No significant group differences in phase entrainment consistency were observed for either the delta (Panel A) or theta (Panel B) bands. However, within both groups, theta-band entrainment exhibited significantly greater consistency than delta-band entrainment.

### 3.4. Comparing band power for the groups across the whole brain

The average power within the analysis time window (see Figure 1) was calculated for each frequency band and group, following the method outlined in Section 2.5. Figure 4 presents boxplots of band-power values for the delta (Figure 4A), theta (Figure 4B), beta (Figure 4C), and low gamma (Figure 4D) bands, separately for the control and dyslexic groups. Based on our hypothesis, we expected group differences in delta- and beta-band powers. To test this, we applied two-tailed Wilcoxon rank-sum tests for each band, excluding outliers. Outliers were defined as power values exceeding 1.5 times the interquartile range above the upper quartile or below the lower quartile within each group. The dyslexic group showed significantly higher power in the delta band than the control group (*z* = –2.43, *p* = 0.015; 2 outliers in the control group), but no significant differences were found for the theta (*z* = –0.94, *p* = 0.35; 2 outliers in the control group, 1 in the dyslexic group), beta (*z* = –0.09, *p* = 0.93), and low-gamma (*z* = 0.01, *p* = 0.99; 1 outlier in the control group, 2 outliers in the dyslexic group) bands.

**Figure 4.**
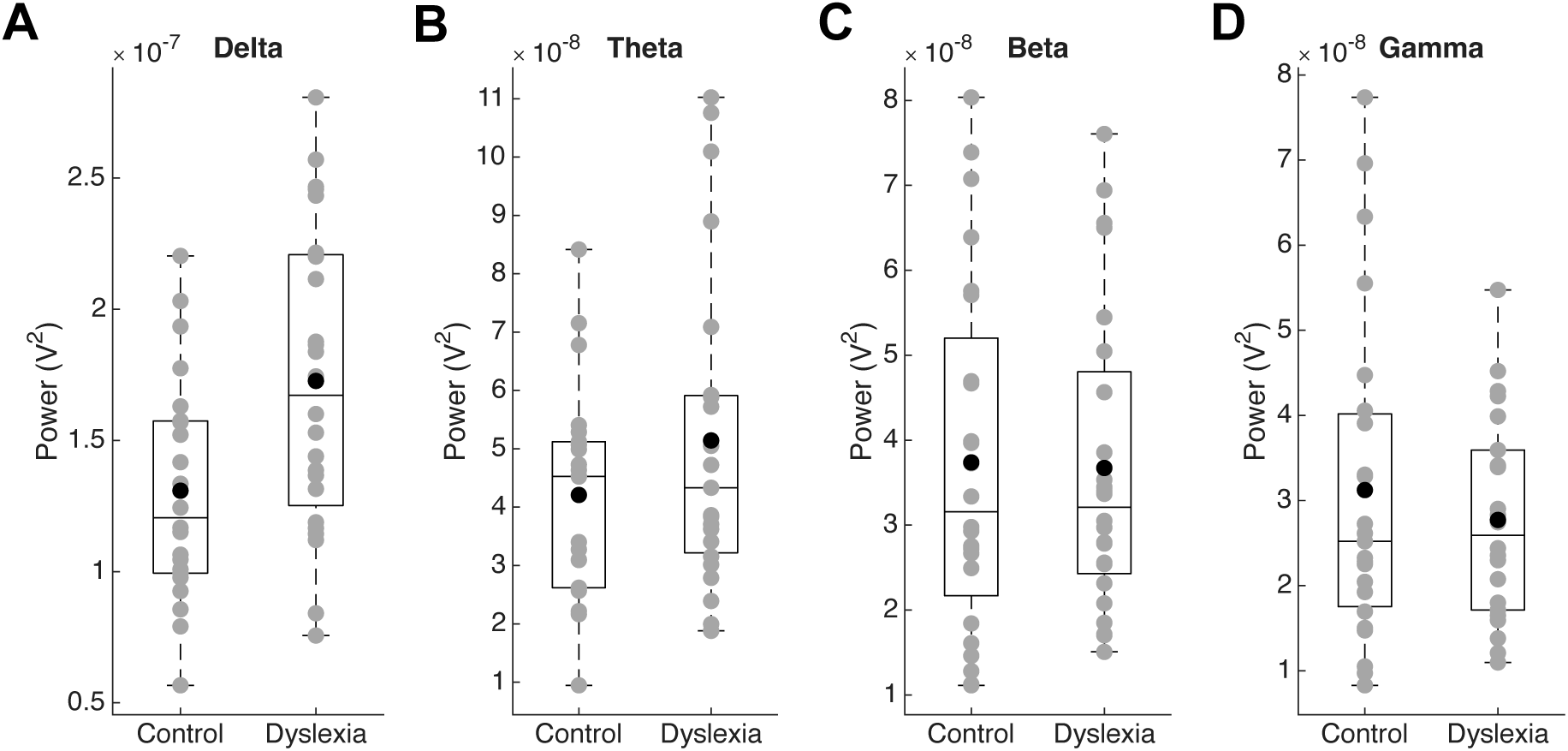
Boxplots of power values for the delta (Panel A), theta (Panel B), beta (Panel C), and low-gamma (Panel D) frequency bands over the analysis time window. Grey circles represent individual participants, black circles indicate group means, and the horizontal line within each box marks the median. Outliers have been excluded from these plots. Note that the y-axis scales differ across panels to reflect the numerical range of power values in each frequency band.

### 3.5. Comparing spectral power topographic maps across groups

Topographic maps were calculated across the entire scalp for each frequency band and group. Figure 5 displays these maps for the delta, theta, beta, low-gamma bands for the control group (Panels A–D), the dyslexic group (Panels E–H), and the difference between groups (Panels I– L). Based on previous findings by Cutini et al. (2016) and Arns et al. (2007), we hypothesised a group difference in delta-band PSD in the right temporal region. To test this, we conducted two-tailed Wilcoxon rank-sum tests for each frequency band within that region. A significant group difference was found for the delta band (*p* = 0.04), but not for the theta (*p* = 0.48), beta (*p* = 0.50) or low-gamma (*p* = 0.32) bands.

**Figure 5.**
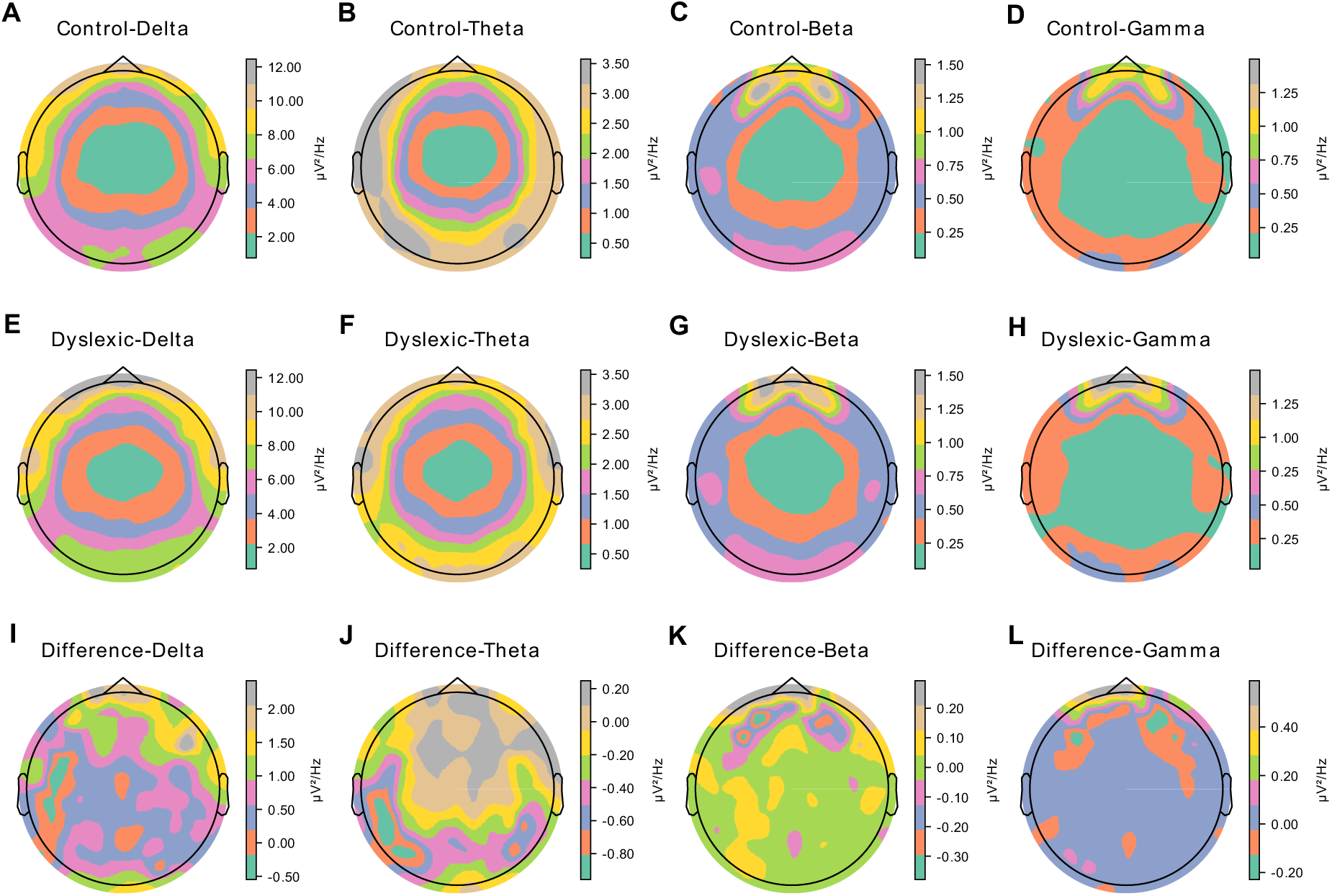
Topographic maps of spectral power for the control and dyslexic groups, and the difference between them. Panels A, B, C, D display the delta, theta, beta, low-gamma band power maps, respectively, for the control group. Panels E, F, G, H show the corresponding maps for the dyslexic group. Panels I, J, K, L present the difference maps (dyslexic minus control) for each frequency band. Colour bars to the right indicate the range of spectral power values; note that the colour scales vary across bands.

### 3.6. Comparing cross-frequency PAC across groups

To examine group-level differences in cross-frequency PAC in response to rhythmic audiovisual stimuli, PAC values were calculated for five frequency pairings: delta-theta, delta-beta, delta-low-gamma, theta-beta, and theta-low-gamma (see Figure S1). These PAC measures were computed separately for the control and dyslexic groups. Group differences in cross-frequency PAC were statistically evaluated using two-tailed Wilcoxon rank-sum tests for each frequency pairing. There were no significant group differences for the delta-theta PAC (*z* = – 1.0, *p* = 0.32; Figure S1A), delta-beta PAC (*z* = –0.69, *p* = 0.49; Figure S1B), theta-beta PAC (*z* = 0.03, *p* = 0.98; Figure S1C), delta-low gamma PAC (*z* = –0.69, *p* = 0.49; Figure S1D), or the theta-low gamma PAC (*z* = 0.13, *p* = 0.89; Figure S1E).

### 3.7. Comparing cross-frequency PPC across groups

PPC values were calculated for the following frequency band pairs: delta-theta, delta-beta, theta-beta, delta-low-gamma, and theta-low-gamma, separately for each group (see Figure S1), using the procedure outlined in Section 2.6. To compare PPC values between groups, two-tailed Wilcoxon rank-sum tests were performed for each band pair. No significant group differences were found for any PPC measure: delta-theta (*z* = –0.18, *p* = 0.86; Figure S1A), delta-beta (*z* = –0.67, *p* = 0.50; Figure S1B), theta-beta (*z* = –0.38, *p* = 0.70; Figure S1C), delta-low gamma (*z* = –0.59, *p* = 0.56; Figure S1D), or theta-low gamma (*z* = –0.09, *p* = 0.92; Figure S1E).

### 3.8. Comparing ERPs across groups

ERP waveforms were computed separately for each group, following the procedure outlined in Section 2.4.9 (see Figure 6). The amplitudes of the N1 and P2 components were compared between groups using separate two-sample t-tests. A significant group difference was observed for the P2 component (*p* = 0.04), while the difference in N1 amplitude did not reach significance (*p* = 0.09).

**Figure 6.**
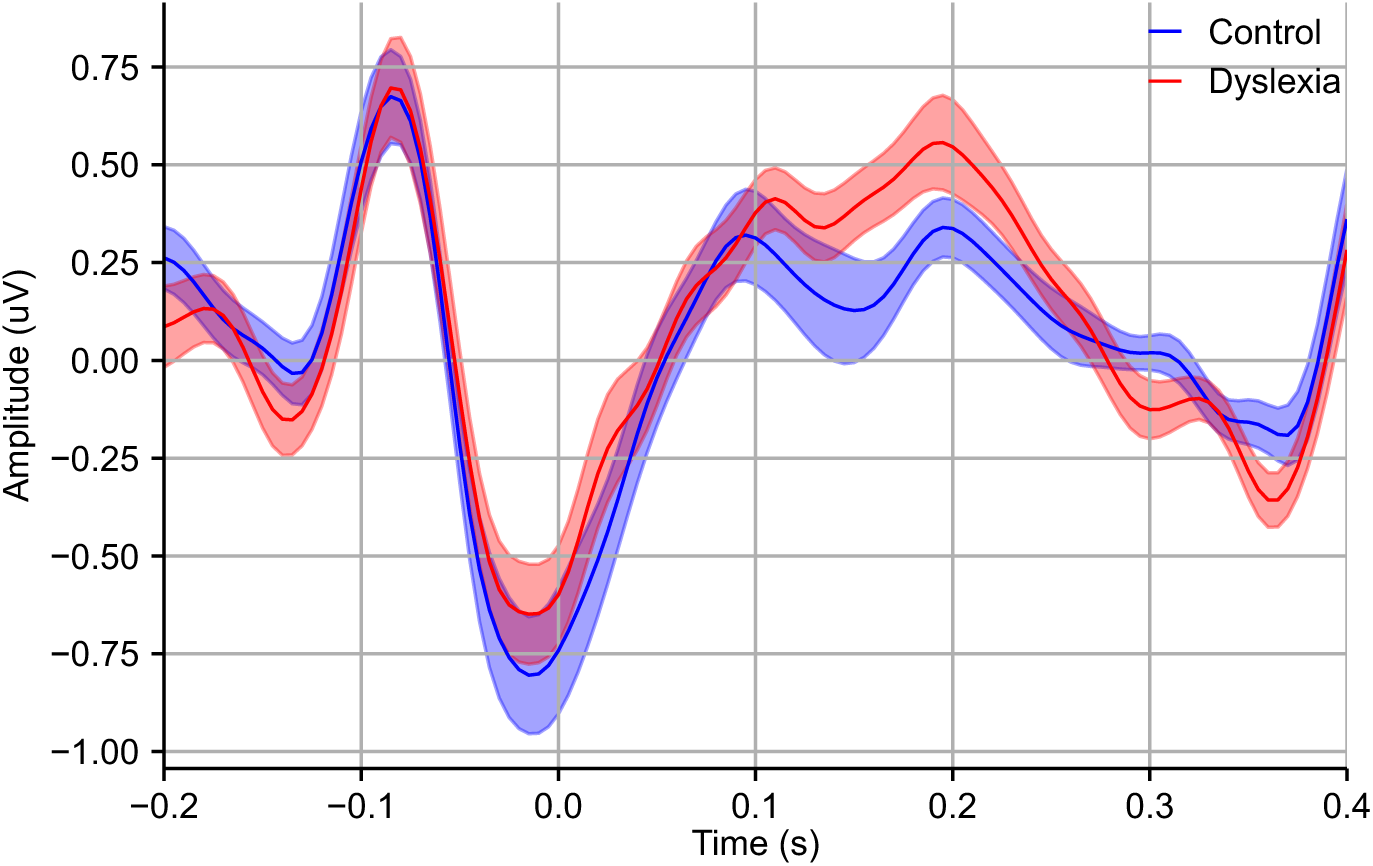
Event-related potentials for control (blue) and dyslexic (red) groups. The shaded areas show the standard error of the mean for each group.

### 3.9. Comparing minimum detectable temporal delays in the audiovisual task (2 Hz) between the two groups

The mean temporal delay thresholds, determined using the adaptive staircase task (see Section 2.2), in which rhythmic sequences of the syllable “ba” were presented with both auditory and visual cues, were 74.3 ms for the control group and 78.5 ms for the dyslexic group. A two-sample t-test showed no significant difference between the groups (*p* = 0.64).

## 4. Discussion

This study was intended to contribute data relevant to the presence of both slow (< 10 Hz, as proposed by TS theory) and fast (beta and low gamma; Lehongre et al., 2011; Poelmans et al., 2012) temporal sampling deficits for adults with dyslexia using rhythmic speech rather than non-speech stimuli, such as AM noise. The study also investigated neural measures of low- and high-frequency cortical dynamics in order to provide a more comprehensive exploration of the neural mechanisms that may contribute to the phonological difficulties observed in individuals with dyslexia (Keshavarzi, 2025). The data revealed significant phase entrainment in the delta and theta bands for both the adult dyslexic and control groups, consistent with previous findings for children with dyslexia (Power et al., 2013; Keshavarzi et al., 2022). The expected group difference in preferred phase of the delta band, as previously reported in studies of children (Power et al., 2013; Keshavarzi et al., 2022), was not found. Instead, the data revealed a significant group difference in the preferred phase in the theta band. As expected from previous TS-driven studies using the rhythmic syllable repetition task or natural speech with children, significant phase entrainment in the beta and gamma bands was absent for the adults with dyslexia. These data are supportive of a low-frequency temporal sampling deficit in individuals with dyslexia, which is next discussed in more detail.

The group difference in preferred phase in the theta band may reflect a temporal delay in neural processing for the individuals with dyslexia. It is possible that, with either maturation, reading experience or both, the phase misalignment characterising the dyslexic brain shifts from the delta to the theta band. For individuals with dyslexia, this phase difference in theta-band activity may reflect impairments in the precise temporal processing of speech. A phase shift in the theta band may also result from the reduced reading experience over developmental time that is associated with dyslexia. In children, increased theta inter-trial coherence in the syllable repetition task is related to greater reading age and to higher standard scores in word and nonword reading (Power et al., 2012). The literate adult brain, with its highly developed orthographic/phonological connections, may hence shift more towards relying on neural mechanisms governed by the theta band during oral speech processing compared to the immature child brain. The atypical preferred phase of dyslexic group observed here for the theta band is consistent with findings from Leong and Goswami (2014), who reported that rhythmic synchronisation at slow rates (∼5 Hz; theta band) was atypical for adults with dyslexia during a behavioural tapping-to-nursery-rhymes task.

Consistent with a maturational account, theta-band phase entrainment consistency was significantly greater than that of delta band consistency for both the dyslexic and control groups. This contrasts with findings for children, for whom no significant difference in phase entrainment consistency has been observed between the delta and theta bands, for either group (Keshavarzi et al., 2022). These results suggest that adults – regardless of dyslexic status – may rely more heavily on theta-band neural dynamics than on delta-band activity during speech processing, at least when processing rhythmic audiovisual stimuli composed of a single syllable. Auditory neuroimaging studies of adults have shown that neural activity in the theta frequency band closely tracks syllable onsets (Ding & Simon, 2014; Di Liberto et al., 2015; Keshavarzi et al., 2020). This contrast between adult and child neural data highlights the importance of considering developmental differences when interpreting neural entrainment in different frequency bands.

Significant phase entrainment was not observed in the beta band for the dyslexic adults, whereas the control group exhibited entrainment for this band. This finding is consistent with our previous results with dyslexic children (Keshavarzi et al., 2024b). The absence of consistent beta-band phase entrainment for adults with dyslexia may indicate a reduced ability to track the timing of temporally regular events. This may reflect deficits in top-down regulatory mechanisms and in temporal prediction – functions typically associated with beta oscillations (Arnal et al., 2015). Indeed, Arnal et al. (2015) demonstrated that beta-band activity is critical for temporal prediction during speech processing in the adult brain, facilitating the alignment of neural oscillations with the timing of upcoming sensory targets, possibly via motor planning (Morillon et al., 2019). Given that Poelmans et al. (2012) also found reduced entrainment for the dyslexic group in the beta band in response to non-speech rhythmic stimulation (AM noise), the absence of consistent beta-band phase entrainment to rhythmic speech stimuli in dyslexia may reflect a difficulty with temporal prediction and sensorimotor integration, which affects both speech and non-speech inputs.

Consistent phase entrainment in the low-gamma band was observed only for the control group. Gamma oscillations have been argued to facilitate the encoding of phonemic elements within speech (Giraud & Poeppel, 2012). Coupled with the data reported by Lehongre et al. (2011) for French adults with dyslexia and AM noise, the EEG data presented here appear to suggest that the gamma band deficits in the French AM noise study are not confined to speech, but instead reflect broader auditory processing difficulties. A subsequent study by Lehongre et al. (2013) using natural speech reported that adult dyslexic readers showed reduced gamma responses in the left hemisphere during passive viewing of an audiovisual movie. These findings were replicated and extended by Lizarazu et al. (2021), who used MEG to demonstrate that dyslexic adults exhibited significantly reduced neural phase-locking at 30 Hz – within the low-gamma range – to both speech and non-speech amplitude-modulated stimuli. Accordingly, the possibility of ‘phonemic oversampling’, which refers to an over-reliance on phoneme-rate temporal information, in adults with dyslexia remains open. It may also be that atypical low-frequency temporal sampling necessarily affects higher frequencies, given the mechanistic hierarchical nesting between slow (delta, theta) and fast (beta, gamma) oscillatory rates in auditory cortex (Gross et al., 2013; Arnal et al., 2015).

Our analysis of ERP components revealed small but significant differences in the P2 component – but not in the N1 component – between the dyslexic and control groups. The N1 component is typically associated with early stages of auditory processing, whereas the P2 component is considered an early neural marker of attentional modulation and timing mechanisms (Hämäläinen et al., 2007), reflecting the allocation of cognitive resources during the perceptual analysis of auditory stimuli. This finding suggests that auditory processing deficits in dyslexia may continue at later cognitive and attentional stages of speech processing, in addition to difficulties associated with the initial sensory encoding of auditory input.

Band-power analysis revealed significant differences between dyslexic and control participants for the delta band, both across the whole brain and in the right temporal region. Differences in the right temporal region may reflect generalised impairments in rhythmic processing, as this region is thought to preferentially process larger units, as reflected in speech and musical prosody (Boemio et al., 2005; Sammler et al., 2015). Both speech and non-speech neural rhythmic studies with children have shown differences in right temporal regions (Arns et al., 2007; Cutini et al., 2016). The absence of significant group differences in the theta and beta bands suggests that the amplitude of oscillations in these frequency ranges may be less affected than oscillations in the delta band for adults with dyslexia, at least in the context of rhythmic audiovisual stimulation.

Our investigation of cross-frequency PPC and PAC revealed no significant differences between the dyslexic and control groups in delta-theta, delta-beta, theta-beta, delta-low gamma or theta-low gamma interactions. These results suggest that synchrony between different neural oscillatory activities – as measured by PPC and PAC – is not significantly disrupted in adults with dyslexia, at least in response to rhythmic audiovisual stimuli. This finding may indicate that neural deficits in dyslexia are more closely related to the neural entrainment and processing of information within individual frequency bands, rather than to cross-frequency interactions. Moreover, the absence of significant group differences in PAC and PPC implies that, although cross-frequency coupling plays an important role in cognition by supporting communication between oscillatory activities in different frequency bands (Canolty & Knight, 2010), it may not be as severely affected in dyslexia as phase entrainment or ERP components.

In conclusion, this study investigated some of the neural mechanisms associated with adult dyslexia by examining neural responses in a rhythmic audio-visual paradigm previously utilised with infants and children. We observed significant phase entrainment in the delta and theta bands for both dyslexic and control groups, with a significantly different theta-band preferred phase for dyslexic individuals, consistent with atypical low-frequency speech processing in dyslexia. Consistent with prior findings with dyslexic children (Keshavarzi et al., 2024b), dyslexic adults lacked consistent beta- and gamma-band phase entrainment. Accordingly, both low-frequency and high-frequency phase entrainment are atypical in the adult dyslexic brain. Both groups of adults exhibited greater theta-band phase entrainment consistency than delta-band entrainment, in contrast to findings with children. This suggests developmental differences that may be linked to either maturation or increased reading experience or both. Finally, significant group differences in delta-band power in the right temporal region may indicate atypical neural oscillatory responses to larger phonological units in speech (such as syllables) in dyslexic adults, consistent with the asymmetric sampling in time proposals of Boemio et al. (2005).

## Author contributions

M.K. conceptualisation, methodology, data collection, data analyses, visualisation, writing-original draft; B.C.J.M. conceptualisation, methodology, writing-review & editing; U.G. funding acquisition, project administration, supervision, conceptualisation, methodology, writing-original draft.

## Acknowledgements

The authors would like to thank all the participants involved in the study. This research was funded by a donation to UG from the Yidan Prize Foundation. The sponsor played no role in the study design, data interpretation nor writing of the report.

## Supplementary Information

**Figure S1.**
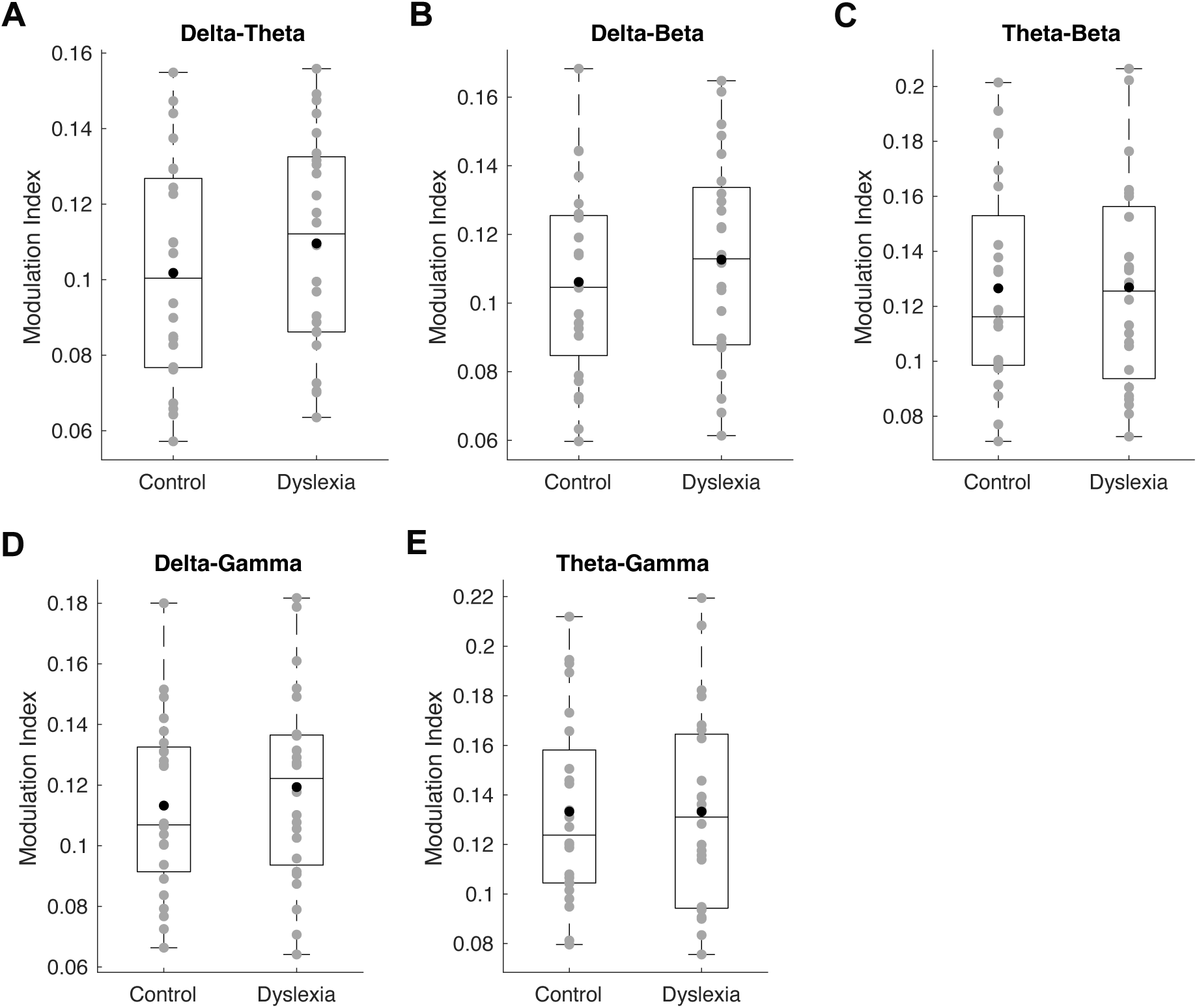
Boxplots of PAC values for each frequency band pair over the analysis time window: delta-theta (Panel A), delta-beta (Panel B), theta-beta (Panel C), delta-low gamma (Panel D), and theta-low gamma (Panel E). Grey circles represent individual participants, black circles indicate the group means, and the horizontal line within each box denotes the median.

**Figure S2.**
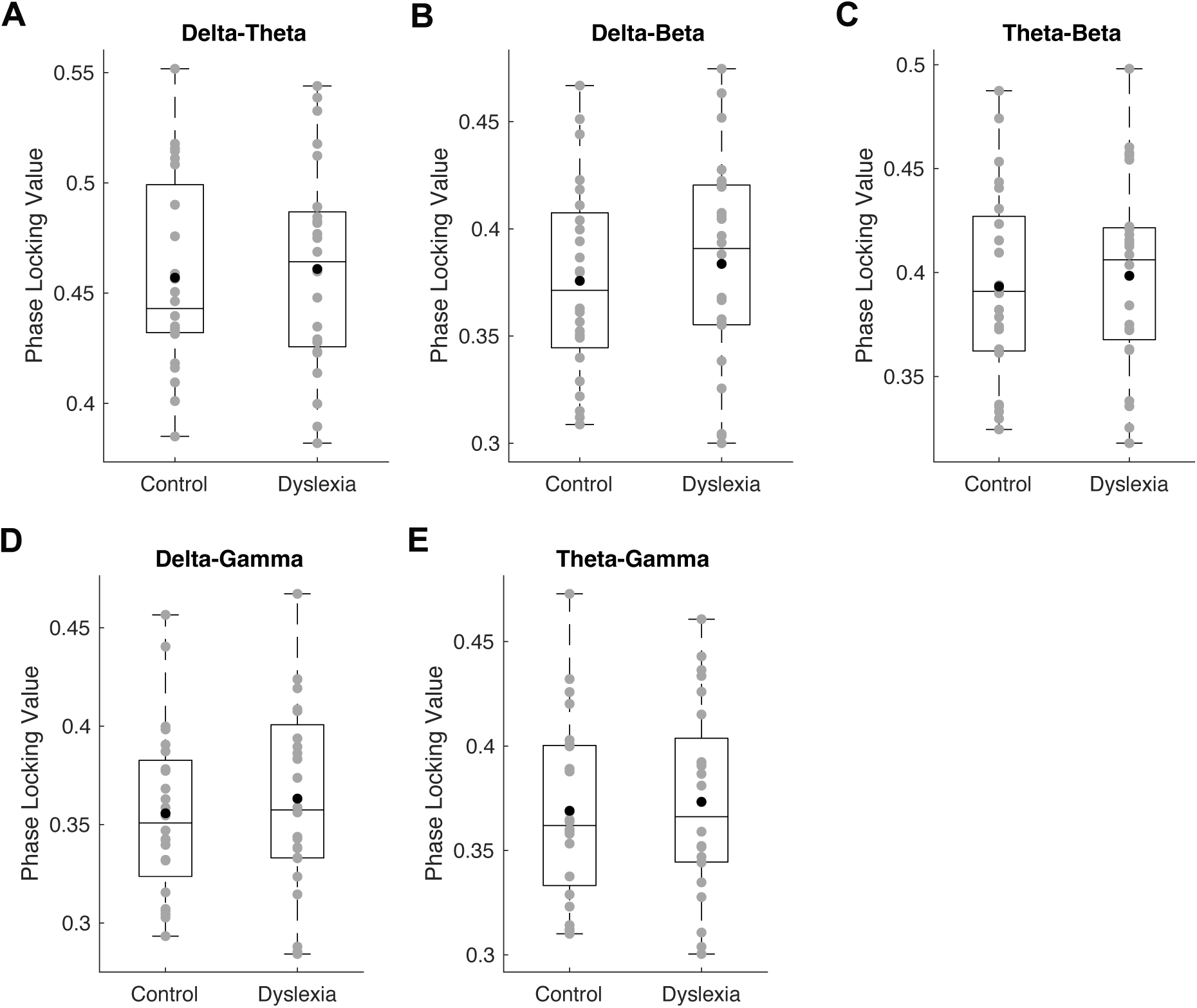
Boxplots of PPC values over the analysis time window for the following frequency band pairs: delta-theta (Panel A), delta-beta (Panel B), theta-beta (Panel C), delta-low gamma (Panel D), and theta-low gamma (Panel E). Grey circles represent individual participant values, black circles indicate group means, and the horizontal line within each box marks the median.

